# Bottlenecks in the Implementation of Genome Scale Metabolic Model Based Designs for Bioproduction from Aromatic Carbon Sources

**DOI:** 10.1101/2024.03.15.585139

**Authors:** Deepanwita Banerjee, Javier Menasalvas, Yan Chen, Jennifer W. Gin, Edward E. K. Baidoo, Christopher J. Petzold, Thomas Eng, Aindrila Mukhopadhyay

## Abstract

Genome scale metabolic models (GSMM) are commonly used to identify gene deletion sets that result in growth coupling, pairing product formation with substrate utilization. While such approaches can improve strain performance beyond levels typically accessible using targeted strain engineering approaches, sustainable feedstocks often pose a challenge for GSMM-based methods due to incomplete underlying metabolic data. Specifically, we address a four-gene deletion design for the lignin-derived non-sugar carbon source, *para*-coumarate, that proved challenging to implement. We examine the performance of the fully implemented design for *p-*coumarate to glutamine, a useful biomanufacturing intermediate. In this study glutamine is then converted to indigoidine, an alternative sustainable pigment and a model heterologous product. Through omics, promoter-variation and growth characterization of a fully implemented gene deletion design, we provide evidence that aromatic catabolism in the completed design is rate-limited by fumarate hydratase activity in the citrate cycle and required careful optimization of the final fumarate hydratase protein (PP_0897) expression to achieve growth and production. A metabolic cross-feeding experiment with the completed design strain also revealed an unanticipated nutrient requirement suggesting additional functions for the fumarate hydratase protein. A double sensitivity analysis confirmed a strict requirement for fumarate hydratase activity in the strain where all genes in the growth coupling design have been implemented. While a complete implementation of the design was achieved, this study highlights the challenge of precisely inactivating metabolic reactions encoded by under-characterized proteins especially in the context of multi-gene edits.

## INTRODUCTION

Computational and data-driven approaches in synthetic biology enable strain performance improvements typically inaccessible using traditional strain engineering approaches. One successful paradigm queries genome scale metabolic models (GSMMs) using growth coupling design algorithms that reroute metabolic flux towards a desired outcome. By implementing the required gene deletions, these changes dramatically improve the productivity of a targeted metabolite or a final product where the chosen metabolite is generated concomitant with microbial growth. Growth coupling has been demonstrated for products using relaxed yield thresholds and model carbon substrates (i.e., glucose) (1,2), but there are fewer examples of such approaches for non-sugar carbon streams (3–5) from renewable bioenergy streams like depolymerized plastics or aromatics from plant-derived lignocellulosic biomass.

The challenge of using growth coupling algorithms for alternative carbon streams is that the existing metabolic data may not fully account for cellular flux outside of glucose (or sugar) catabolism, where flux information is less robust. If the modeling does not reflect true cellular metabolism, these approaches are more likely to generate gene deletion targets that are ineffective or inaccurate. Growth coupling algorithms, which promise substantial improvements to titer rates and yield (TRY) are termed “strong” growth coupling, also demand many gene modifications (upwards of 15 gene deletions) that are nontrivial for implementation (6). Both partial and complete implementation of such designs, by themselves or in conjunction with other approaches have shown useful applications for bioproduction (1,7–9) and reveal key insights into the host strain physiology.

We use a previously developed strong growth coupling design in *Pseudomonas putida* KT2440 that pairs a lignin-derived monomer, *para-*coumarate (*p*-CA) to glutamine (10) (**Figure 1**), a valuable platform chemical (11–13). We further use heterologous expression of a non-ribosomal peptide, indigoidine, an alternative for industrial petrochemically-derived indigo (14–16). Indigoidine is generated by condensation of two glutamine molecules (**Figure 1**) and is often used as a proxy for glutamine, measured using quantitative colorimetric assays for efficient strain prototyping (15,17,18). Non-model carbon sources pose a challenge for growth coupled strategies, as substrate toxicity can be a challenging baseline for strain engineering (19). We used <u>c</u>onstrained <u>m</u>inimal <u>c</u>ut<u>s</u>ets (cMCS), a computational approach (1,6) that provides strong coupling solution-sets or cut-sets, where each cutset consists of reactions (and the corresponding genes) that need to be deleted for the growth coupled production phenotype. For bioconversion to glutamine/indigoidine, we showed that a partially implemented cutset (three out of the four demanded genes were deleted) enabled phenotypic growth coupling, but generated the product at lower levels than the predicted yield (10). As we encountered unexpected strain behavior after three of the four deletions were introduced, we used an ensemble of methods to characterize the triple deletion strain. However, since an implementation of the complete cutset was not accomplished, a hypothesis that emerged is that a full cutset may allow higher productivity as predicted for the strong growth coupling design without additional refinement, or provide key insights into the role of the reactions involved in this design.

**Figure 1.**
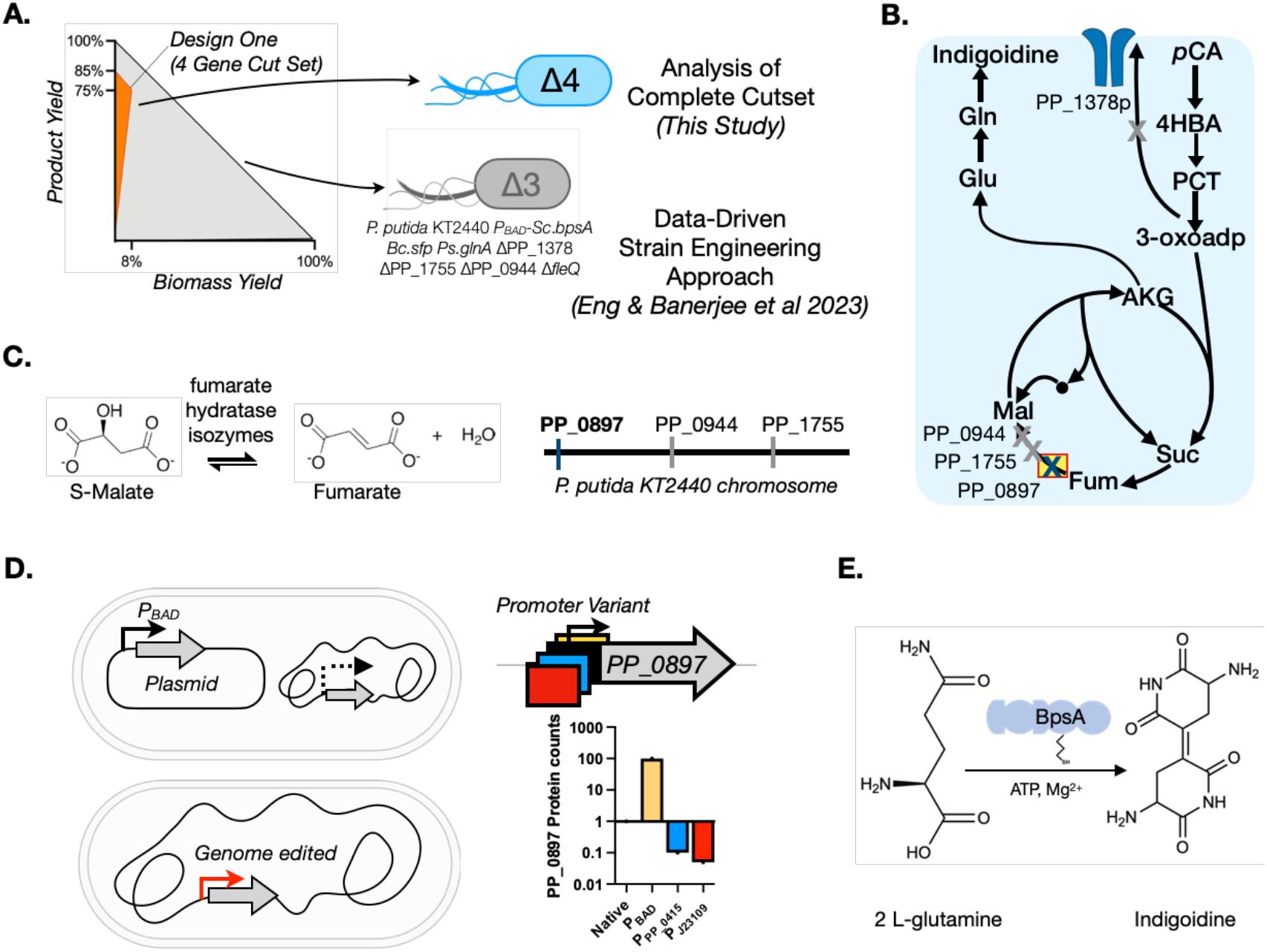
Growth Coupled Cutset for the Bioconversion of *para*-Coumarate to glutamine and Indigoidine. (A) Yield envelope for this study. The complete growth coupling solution described with the orange fill area (Design 1b) has a predicted MTY of 76% indigoidine from *p-*CA. The yield is shown for a range of substrate uptake rates from 0 - 10 units. For comparison, the predicted biomass formation rate and yields for a partial cutset (Eng & Banerjee et al 2023) are included for comparison as the grey area fill in the yield envelope. (B) Simplified metabolic map showing the four required gene interventions necessary to implement the Design 1b cutset in *P. putida*. Three genes involved in fumarase hydratase activity and one permease must be deleted. (C) The reaction and genomic locations of the three genes involved in fumarase hydratase activity. (D) The different approaches taken to modulate the expression of PP_0897, the last fumarate hydratase gene, that needs to be deleted for a complete cutset implementation. (E) Schematic representation of the indigoidine production module using 2 molecules of L-glutamine converted to the blue pigment, a non-ribosomal peptide.

Here, we characterize the behavior of an engineered *P. putida* strain containing the complete cutset and the impact on heterologous final product formation. We demonstrate that the fourth gene in this cutset, PP_0897, is involved in cellular activity beyond its annotation as a fumarate hydratase. Overexpression of PP_0897 decreases final product titer and growth, while its deletion leads to unexpected auxotrophies. Constraint-based analysis of context-specific models, using proteomics data, was in good agreement with these experimental observations. The analysis of the complete cutset identifies a new bottleneck in the citrate cycle governing *p*-CA catabolism and in turn its conversion to potentially other products. A double sensitivity analysis (20,21) demonstrates a strict requirement for fumarate hydratase activity and the need to validate growth coupling predictions with additional cellular information, potentially applicable for other glutamine-derived bioproducts.

## RESULTS

### A Completed *p*-CA/Indigoidine Cutset and Modulating the PP_0897 Gene Impacts Strain Growth, Production and Proteome Response

The complete intervention design (cutset) requires deletion of four genes: an α-ketoglutarate / 3-oxoadipate permease PP_1378; and three class I and II fumarate hydratases - PP_0897, *fumC1*/PP_0944, and *fumC2*/PP_1755. This design was reported in Eng et al 2023 (10) where a partial implementation deleting only PP_1378, PP_0944, and PP_1755 was used and the corresponding strain was further engineered using data driven approaches for production improvement (**Figure 1A** and **1B**). The conceptual foundation from this design was an appealing starting point to test our initial hypothesis. By including one additional deletion (PP_0897) we could complete the cutset, potentially revealing growth coupling at the predicted product yield without additional model refinement. We started with the strain *P. putida* KT2440 *P*_*BAD*_-Sc.*bpsA*, Bc.*sfp*, Ps.*glnA* ΔPP_1378 ΔPP_1755 ΔPP_0944 *ΔfleQ (10), i*.*e*. Design 1b *glnA* Δ*fleQ* (see strain table, **Supplementary Table 1**, abbreviated and hereafter referred to as “D1b_gf”). The fumarate hydratases in the design are associated with the FUM reaction in the *P. putida* GSMM (**Figure 1B** and **1C**). Deletion of either or both the non-essential fumarate hydratases, PP_1755 and PP_0944 had no impact on cell viability in either rich or M9 *p-*CA minimal media cultivation (10).

We tested if PP_0897 was an essential gene in *P. putida* by targeting this gene alone for deletion via CRISPR/recombineering. PP_0897 deletion mutants were recovered at a comparable frequency expected for our established recombineering protocol (Materials and Methods). *P. putida* ΔPP_0897 cells are viable on both rich and *p*-CA minimal salt medium but form smaller, slower growing colonies (**Figure 2A**). These results indicate that PP_0897 alone is dispensable for growth in both rich and M9 *p-*CA minimal medium conditions. However, D1b_gf ΔPP_0897 strains (containing all 4 deletions) were slow growing on LB medium and failed to grow on M9 *p-*CA medium agar plates (**Figure 2A**) and also did not produce any final product (**Figure 2C)**. To test if serial passaging would restore both growth and production of a completed cutset, we subjected D1b_gf ΔPP_0897 to adaptive laboratory evolution. We failed to recover any improved mutant clones after three months of serial passaging on rich or minimal medium, at which point the experiment was discontinued.

**Figure 2.**
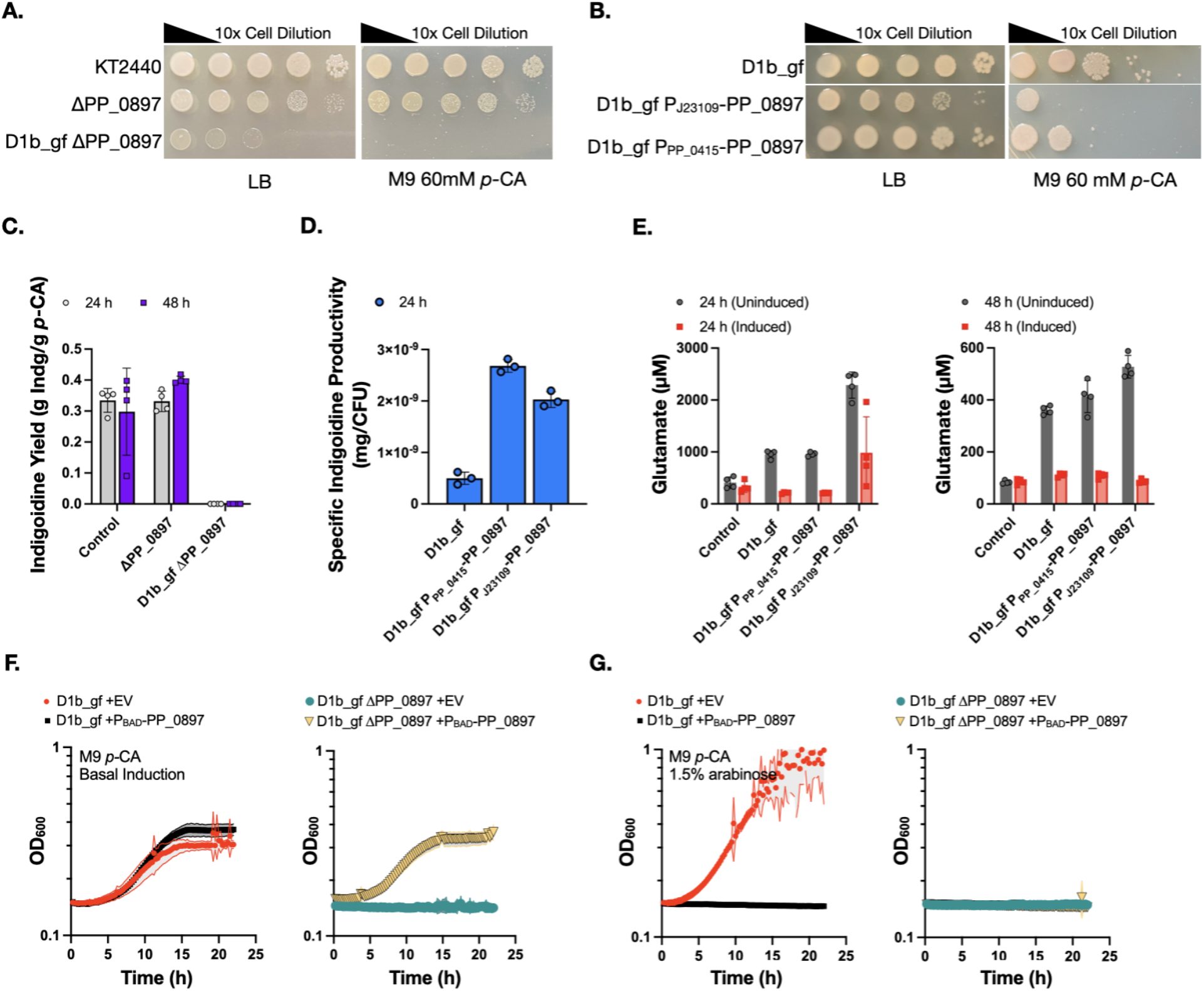
Relationship between the PP_0897, fumarate hydratase expression, growth and production. (A) Serial dilution plating for growth on solid agar medium supplemented with rich (LB) or *p*-CA minimal medium. Abbreviated genotypes are indicated to the left of each image. (B) Native PP_0897 promoter was replaced with two different low abundance promoters and the impact on viability in rich medium or M9 *p-*CA minimal medium was quantified. (C) *P. putida* strains assayed for indigoidine production in M9 60 mM *p*-CA medium at the indicated time points following induction with 1.5% arabinose (w/v). Control is WT *P. putida* harboring genome integrated indigoidine pathway cassette and ΔPP_0897 is the single gene deletion control. Strain D1b_gf ΔPP_0897 did not grow or produce in M9 *p-*CA medium. (D) *P. putida* strains with promoter varied PP_0897 expression assayed for Indigoidine production. Specific productivity (mg/CFU) was calculated by normalizing the indigoidine with the number of total viable cells at 24 hrs. (E) *P. putida* strains with promoter varied PP_0897 expression with or without induction of the indigoidine pathway in M9 60 mM *p-*CA, sampled at 24 h and 48 h for glutamate and indigoidine (Supplementary Figure 1). (F and G) Strains D1b_gf with or without ΔPP_0897 were transformed with an empty vector plasmid or a plasmid containing a *P*_*BAD*_-PP_0897 construct. (F) Growth in M9 60 mM *p-*CA medium with basal *BAD* promoter induction without inducer or with 1.5% (w/v) arabinose (G) in 24 well microtiter dishes. Strain genotypes are indicated above each panel. Measurements are average of at least three independent replicates. Error bars represent mean ± S.D. (n=4) in C, D and E. Shaded area represents mean ± S.D. (n=3) in F and G.

Since the cutset could not be completed by deleting PP_0897 due to strain inviability (**Figure 2A)**, we asked if a reduced PP_0897 expression strain would permit growth and glutamine/indigoidine production with *p-*CA as the carbon source. The endogenous PP_0897 promoter was replaced using recombineering with either a low activity pJ23109 promoter from the Anderson collection (22) or the PP_0415 promoter sequence, that in previous proteomics characterization had been established to be a low abundance protein in the D1b_gf strain (10) (**Figure 1D**). While D1b_gf P_PP_0415_-PP_0897 had a modest growth reduction compared to the base engineered strain (WT harboring an integrated indigoidine production pathway), strain D1b_gf P_J23109_-PP_0897 required an additional 24 h incubation on LB agar medium to form colonies (**Figure 2B)**. We observed colony size heterogeneity in the D1b_gf P_J23109_-PP_0897 strain; sanger sequencing of the J23109 promoter indicated that the larger and faster growing clones accumulated a G/A point mutation in the -22 position upstream of the PP_0897 start codon. Passaging the smaller colonies invariably led to restored growth rates and the appearance of the same -22 position G/A point mutation, suggesting the chosen Anderson promoter expression level was below a threshold necessary for growth in rich medium growth conditions that favored spontaneous accumulation of the point mutant. In contrast to D1b_gf ΔPP_0897 strains, both of these PP_0897 promoter variants enabled growth on M9 *p*-CA **(Figure 2B)** without serial passaging, providing us with a route to investigate the role of this gene further.

Next we examined the impact of the various PP_0897 edited configurations on the accumulation of the target metabolite, glutamine and on indigoidine production. We observed that the single deletion strain, ΔPP_0897, showed similar product yields to the basal pathway engineered strain while D1b_gf ΔPP_0897 did not show any production (**Figure 2C**). In contrast, the reduced expression promoter mutants accumulated similar amounts of indigoidine to the parental control strain (**Supplementary Figure 1**). By accounting for the number of viable cells present during a production run, we observed that the specific indigoidine productivity in the promoter variants was up to 5-fold higher compared to the parent D1b_gf strain (**Figure 2D**). These results indicate that native PP_0897 activity is necessary for robust biomass formation as well as conversion of glutamine to indigoidine. The promising per cell increase in indigoidine led us to further investigate the lack of growth on the defined *p*-CA medium. Since the growth-coupling design targets glutamine, we measured the glutamine and glutamate pools via LC-MS, but the glutamine concentrations were below the detection limit for all samples in this study. Consistent with growth coupling, the promoter varied strains showed increased glutamate pools (**Figure 2E**). The strain D1b_gf P_J23109_-PP_0897 had the highest glutamate pool increase of 2.4-fold relative to the control strain. Induction of the indigoidine production pathway depleted the glutamate pools in all three growth-coupled strains. In these cases *glnA* is also induced as part of the optimized heterologous pathway used in the D1b derived strains. In contrast, the control strain showed no detectable change in glutamate levels as it lacks an additional, inducible copy of *glnA* (**Figure 2E**). However, the increased glutamate concentration did not correlate with additional indigoidine, and indigoidine yields were similar across strains (**Supplementary Figure 1**). The cap on improvement in indigoidine could be reflective of insufficient cofactors to complete the cyclization of glutamine by BpsA, limiting additional conversion of this intermediate. Additional evaluation of redox balance for these strains are required to confirm this hypothesis. The HPLC analysis confirmed that the PP_0897 promoter variants consumed all of the supplied *p*-CA within 24 h post inoculation. These results indicate that native PP_0897 activity is necessary for robust biomass formation, the growth coupled conversion to glutamine, and for the generation of indigoidine.

To rule out the impact of secondary mutations in the D1b_gf strain background on the failure to grow in M9 *p-*CA medium when PP_0897 is deleted, we designed new plasmids to reintroduce PP_0897 under an inducible promoter. This allowed us to investigate the association of PP_0897 and the synthetic lethal phenotype observed in D1b_gf ΔPP_0897. D1b_gf and D1b_gf ΔPP_0897 strains were transformed with either an empty vector plasmid or an arabinose inducible *P*_*BAD*_-PP_0897 construct. We observed that the *P*_*BAD*_-PP_0897 restored growth of D1b_gf ΔPP_0897 strains with basal induction (**Figure 2F**), confirming genetic linkage by complementation. However, inducing PP_0897 with higher arabinose concentrations (1.5% w/v) inhibited strain growth (**Figure 2G**), revealing a cellular sensitivity to dosage level. Over-expression toxicity was not limited to the D1b_gf ΔPP_0897 strain but also to the parental D1b_gf strain, and the basal expression of PP_0897 restored production in D1b_gf ΔPP_0897, but reduced the final titers of D1b_gf by 3.8-fold (**Supplementary Figure 2A**). In summary, these results suggest that the accuracy of a functioning complete cut-set is impacted by gene essentiality, as well as secondary metabolic interactions resulting from multiple deletions.

To characterize global perturbations on cell metabolism from reduced PP_0897 protein levels, we used proteomics to identify differentially expressed proteins. We found 91 and 56 proteins that were respectively significantly up or down regulated in the two promoter variant strains (P_J23109_ and P_PP_0415_) based on the log_2_ fold changes compared to the parental D1b_gf strain (**Supplementary Figure 3A**). Out of this set, we focused on the subset of proteins that were common between both promoter mutants to set a high threshold for potential biological relevance. Out of the ∼2,000 quantified proteins captured by proteomics, there were only 28 metabolic and 63 non-metabolic differentially expressed proteins, in the P_J23109_ strain. Similarly, the P_PP_0415_ strain contained only 19 metabolic and 37 non-metabolic differentially expressed proteins. Only 13 proteins were common between both strains. Subsequent proteomics analysis confirmed that the PP_0897 protein was enriched 120-fold over PP_0897 protein counts observed in the D1b_gf with the empty vector control (**Supplementary Figure 2D**). Shotgun proteomics analysis confirmed PP_0897 abundance in the promoter variant strains were reduced in the D1b_gf P_J23109_ strain by log_2_ fold of -4.04 and by log_2_ fold of -3.28 in the D1b_gf P_PP_0415_ strain (**Supplementary Figure 3B**). Basal PP_0897 expression in the D1b_gf ΔPP_0897 with the *P*_*BAD*_-PP_0897 construct was already 5-fold higher than the same reference strain, indicating modest increases in expression level were tolerated. Besides the expected deleted gene PP_0897, other metabolic proteins including VanA, PP_5387; transport related proteins AapJ, PP_4379, and PP_2797; and putative uncharacterized proteins PP_2575 and PP_5304 were also downregulated. Barcoded transposon fitness information for these 13 genes (**Supplementary Figure 3C**) indicates that inactivating most of these proteins as single mutants has limited impact on cellular fitness when *P. putida* is grown on *p-*CA or related aromatic compounds with the exception of VanA, which has a defect only on vanillate but not *p-*CA.

### Context Specific GSMM for Strains with Various Growth Coupling Implementation Explains the Highly Constrained Design Space

Our empirical evidence suggested that modulating PP_0897 activity was not reflected in the GSMM based designs for *p*-CA conversion to glutamine. Thus, we hypothesized that the metabolic model and cMCS based high yielding predictions for growth coupled strategies were over-constrained under experimental scenarios using *p*-CA and there may be metabolic and non-metabolic processes that are unaccounted for. One solution is to refine the models to account for complex multilayered networks for regulation using functional genomics data as well as regulatory network information (23). Here, we specifically used context specific models generated using proteomics data for the two promoter-based strains as an alternative approach/solution.

Our analysis confirms a narrow range of permissible flux through the FUM reaction that would result in strong coupled glutamine production, *i*.*e*. a fixed amount of glutamine flux is required at all times while generating biomass for the engineered strains when utilizing *p-*CA as the sole carbon source (**Figure 3**). We found this by utilizing differential proteomics data as new constraints for the D1b_gf reduced model (iMATD1b539) which has 558 reactions and 539 genes (10). This generated two different context specific models for the P_J23109_ strain (iMAT_pJ23109_PP_0897, 522 reactions) and the P_PP_0415_ strain (iMAT_PP_0415p_PP_0897, 522 reactions) (**Figure 3A**, refer to Proteomics data integration to create context specific models section in Methods). Proteomics data analysis for the two promoter-based variants, P_J23109_ strain and P_PP_0415_ strain showed that there were 27 and 18 proteins, respectively, that had a significant differential protein abundance compared to D1b_gf strain. Based on previous data (10) we were able to derive growth, glutamine and indigoidine flux for the two context specific models. The predicted maximum biomass was 1.14 hr^-1^ and the maximum glutamine flux was 8.77 mmol/gDCW/hr (i.e. 0.88 mol/mol) using *p*-CA as the sole carbon source. Next, using flux variability analysis and robustness analysis we observed that the balance between glutamine and biomass flux was highly dependent on the fumarase (FUM) reaction flux for the promoter-based strain specific models compared to the GSMM as well as the D1b_gf reduced model (**Figure 3 B - D**). For maximal glutamine production, FUM carried a flux of 19.88 mmol/gDCW/hr which resulted in 0.43 hr^-1^ biomass whereas for maximal biomass, the FUM reaction carried 26.44 mmol/gDCW/hr flux that resulted in a negligible glutamine flux. Increasing the FUM flux by a small fraction resulted in growth coupled production phenotype i.e. constraining the FUM flux to 20 mmol/gDCW/hr results in 0.45 hr^-1^ biomass led to a coupled production of 8.61 mmol/gDCW/hr (i.e. 0.86 mol/mol) glutamine. This suggests a potential tradeoff between maximal growth and production and also that a nominal change in FUM flux under the defined medium conditions and proteomics constraints could result in the desired growth coupled phenotype.

**Figure 3:**
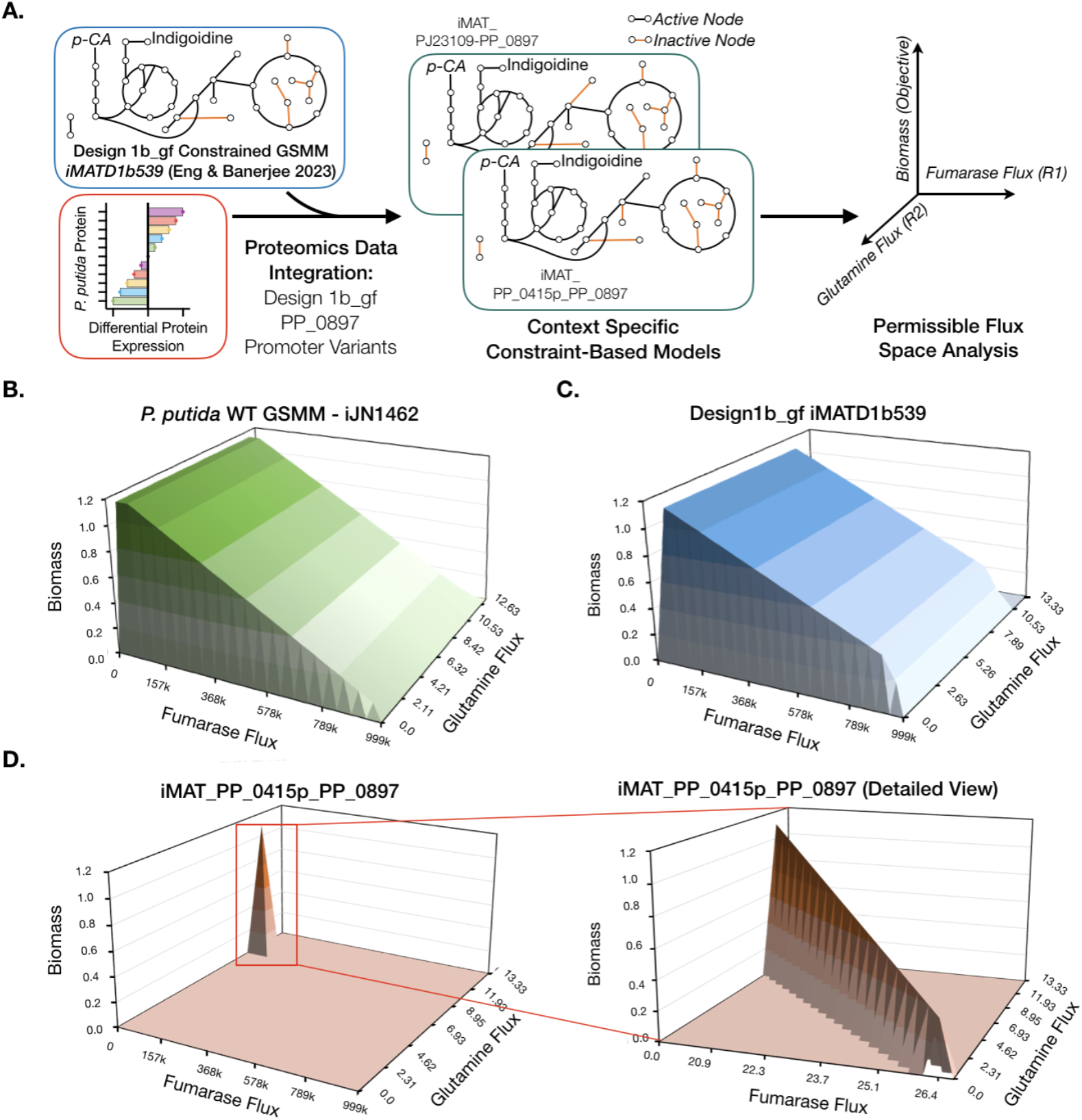
Proteomics-constrained context-specific metabolic models reveal narrow permissible fumarase flux space for growth coupled production. (A) Overall workflow of the simulations performed to model the permissible flux space limiting growth coupled production. Proteomics data, D1b_gf reduced model and the iMAT algorithm are used to develop context-specific models which are then used for sensitivity analysis. A double robustness analysis is applied to determine the tradeoff between growth, fumarase flux and glutamine/indigoidine formation. Double robustness analysis 3D plots of the genome scale metabolic model (GSMM) of *P. putida* WT (iJN1462), (B), context specific D1b_gf reduced model (C) and context specific D1b_gf P_PP_0415_-PP_0897 model (iMAT_PP_0415p_PP_0897). A red box indicates the region of interest replotted in the detailed view.

To assess the interplay between all three parameters; growth, the FUM reaction, and magnitude of production, in the entire available flux space for the context specific models, we performed double sensitivity analysis (via double robustness simulation, see Methods). In contrast to the unconstrained GSMM and the D1B_gf model, double sensitivity analysis predicted a single point FUM flux (**Figure 3D**) that allowed strict growth coupled production of glutamine. Both the context specific models for the promoter-based strains i.e. iMAT_pJ23109_PP_0897 and iMAT_PP_0415p_PP_0897 showed similar trends. As expected for the growth coupled design, we observed similar balance and narrow permissible FUM flux trends for indigoidine production (**Supplementary Figure 4**). The narrow flux peak visualized could explain the failure to recover ALE strains for D1b_gf ΔPP_0897 because this flux may be biologically infeasible with the demand for both reasonable biomass formation and coupled production with *p*-CA as the sole carbon source.

### Complete Implementation of All Four Gene Deletions Results in Multiple Auxotrophies

The computational and experimental evidence indicated that the complete cutset strain had stringent requirements for growth on *p-*CA and suggested the carbon source was rate limiting for biomass formation. A simple explanation was that malate, the product of fumarate dehydratase, was necessary for growth due to the complete inactivation of that TCA node in the D1b_gf ΔPP_0897 strain. D1b_gf ΔPP_0897 failed to grow in M9 60 mM *p-*CA with malate supplementation (**Figure 4A**) in contrast to the predicted growth from the constraint-based modeling of a promoter titrated strain and analysis. Both of these growth trends contrast with the complementation analysis, where the PP_0897 coding sequence introduced on a plasmid was sufficient to restore growth in ΔPP_0897 strains in both M9 *p*-CA (**Figure 2F**) and M9 *p-*CA malate (**Figure 4A**), suggesting a malate auxotrophy did not explain the observed growth defect in ΔPP_0897 strains. Malate supported growth as the sole carbon source in control strains, indicating malate transport was not a limiting factor **(Supplementary Figure 5**).

**Figure 4:**
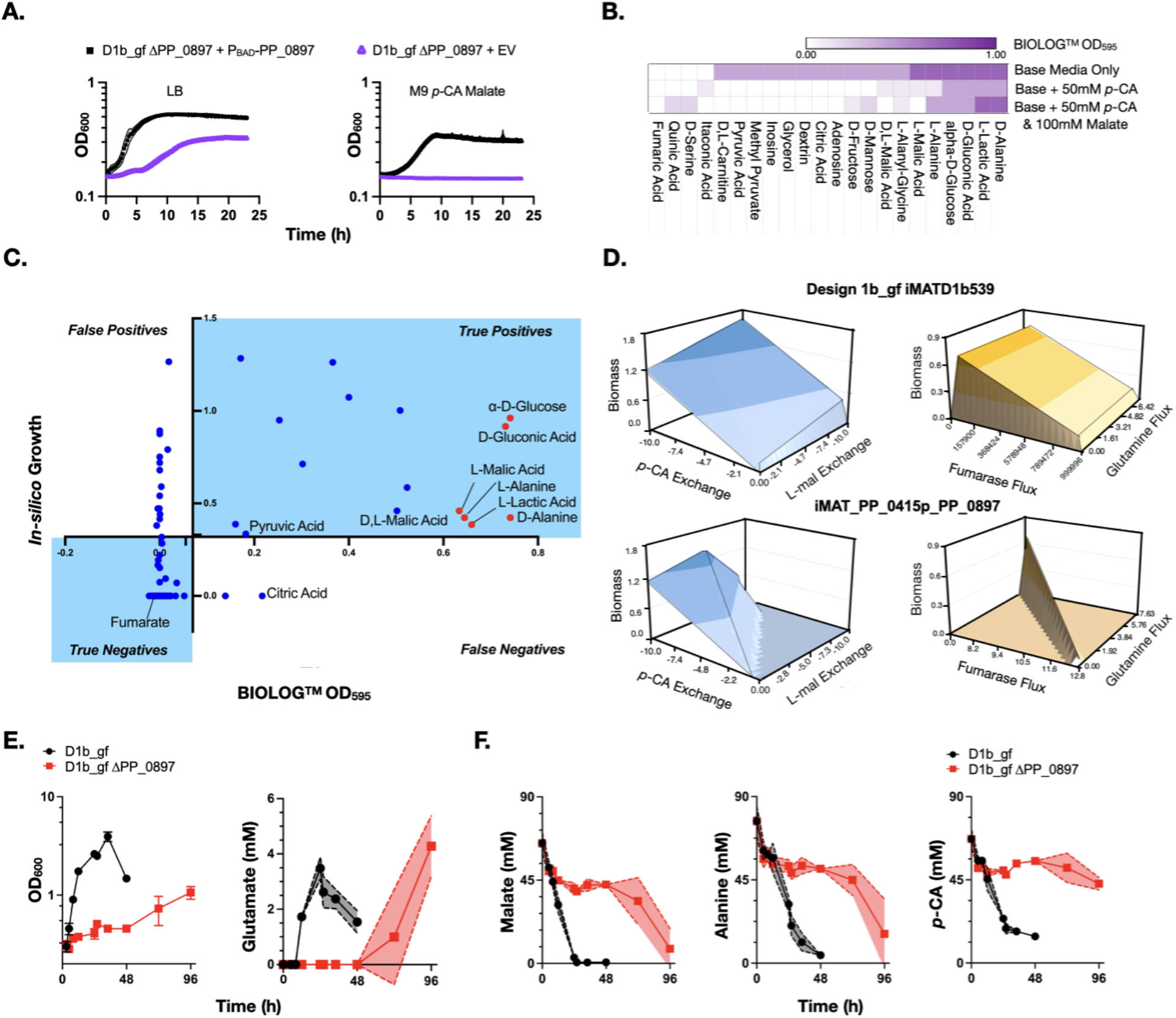
Metabolite Supplementation to Rescue Growth of D1b_gf ΔPP_0897. (A) Initial auxotrophy analysis of the completed Design 1 cutset, D1b_gf ΔPP_0897 harboring PP_0897 on a plasmid. (B) Heatmap showing selected BIOLOG™ metabolite sources that rescued growth under different media conditions including basal media supplemented with 50 mM *p*-CA and/or 100 mM L-malate. (C) Model validation using BIOLOG™ phenotyping. Blue dots indicate each of the 190 metabolites tested (Refer Supplementary Table 4 for details), light blue shaded region shows the accurately predicted metabolites including 16 true positives and 75 true negatives. Red dots indicate the subset of metabolites (6) that successfully rescued growth fed both *p*-CA and L-malate. (D) Double robustness analysis on mixed carbon source utilization for growth and production. (E - G) Strains D1b_gf and D1b_gf ΔPP_0897 were grown using three carbon sources, 50 mM *p*-CA, 100 mM malate, and 100 mM alanine and culture samples were taken at the indicated time points for growth and metabolic profiles. E) *Left:* Optical density (OD_600_) was monitored every 4 h. *Right:* Glutamate concentrations for strains D1b_gf and D1b_gf ΔPP_0897. F) Consumption profiles of the three carbon sources malate, alanine and *p*-CA monitored at the indicated time points using LC-MS. Error bars represent mean ± S.D. (n=3) in E (left). Shaded region represents mean ± S.D. (n=3) in A, E (right) and F.

To pin-point other potential metabolites that would enhance growth on *p-*CA, we considered media amendments that could supplement *p-*CA as the sole carbon source. Since malate supplementation did not restore D1b_gf ΔPP_0897 growth in M9 *p-*CA, we searched for additional missing nutrients using BIOLOG™ phenotyping microarray plates. These plates contain 190 metabolites arrayed in two 96 well microtiter plates enabling rapid nutrient utilization profiling. *P. putida* KT2440 was reported to utilize 47 of 95 of the metabolites in PM1 plate (24), while a “Δ flagella” strain lost the phenotype to utilize D- or L-alanine (25). In contrast, our strain D1b_gf ΔPP_0897 was able to utilize both alanine isomers, even though it contains a comparable Δ*fleQ* edit, inactivating the master regulator for flagella biosynthesis. D1b_gf ΔPP_0897 lost the utilization phenotype for 20 metabolites and resulted in the highest signal of cellular respiration with an OD_595_ above 0.7 for six metabolites - D-glucose, D-alanine, L-alanine, L-malic acid, D-lactic acid and D-gluconic acid (**Figure 4B**).

Next, we cross-validated the growth phenotypic data against the GSMM using FBA to determine model fidelity for substrate utilization in a completed Design 1 strain. For the PM1 BIOLOG™ phenotype plate, 80 out of 95 substrates tested were predicted accurately using the GSMM, with an accuracy of 84% (**Figure 4C**). In the case of the PM2A plate, 11 were predicted accurately using the GSMM including biomass formation using L-carnitine for the D1b_gf ΔPP_0897 strain. Overall, for the two BIOLOG™ plates, with only minor modifications to the existing GSMM, an overall 78% prediction accuracy indicates the robustness of the *P. putida* GSMM for predicting substrate utilization in a growth coupled strain. However, the addition of *p-*CA to the base medium has a dominant negative impact on the predictive modeling accuracy. When the baseline medium was supplemented with 50 mM *p*-CA, fewer substrates were able to drive respiration with *p-*CA supplementation, reducing the number of candidate supplemental metabolites from 18 to 6, including a-D-glucose, D-gluconic acid, L-malic acid, L-alanine, D-alanine and L-lactic acid (**Figure 4C**). The strongest growth benefits were observed for D-alanine and L-lactic acid, two unintuitive metabolites distal from the TCA cycle. Using double robustness analysis we also confirm that for both the promoter based strains there exists an even more constrained permissible flux space for growth and production when *p*-CA was supplemented with malate, using the context specific iMAT_PP_0415p_PP_0897 model (**Figure 4D** compared to **Figure 3D**). As we had hypothesized that there would be a second auxotrophy in addition to a malate auxotrophy, we identified media conditions that restored growth in the presence of both *p-*CA and malate. Fumarate supplementation was never detected as a growth-enhancing metabolite, in contrast to some previous reports in *Arabidopsis thaliana* (26) that fumarate dehydratase can catalyze both the forward and reverse reactions.

Next, we selected D-alanine and L-malate to test whether they would rescue both metabolic respiration and growth in the strains D1b_gf and D1b_gf ΔPP_0897 in the presence of *p-*CA. A combination of 50 mM *p*-CA, 70 mM L-malate and 70 mM D-alanine worked best for the strain D1b_gf ΔPP_0897 but growth was only partially restored, reaching an OD_600_ of 1.0 after 72 h of cultivation **(Figure 4E)**. Metabolite analyses revealed that the growth of D1b_gf ΔPP_0897 was mainly supported by the co-consumption of L-malate and D-alanine, while *p*-CA use was limited **(Figure 4F)**. In contrast, the growth of the parental strain D1b_gf was robust and this strain co-consumed all three carbon sources by 48 h. Further, in D1b_gf despite metabolizing the wider set of available carbon sources, indigoidine production was enhanced only 1.2 times more compared to M9 60 mM *p*-CA alone (**Supplementary Figure 6**). D1b_gf ΔPP_0897 produced 10-fold lower indigoidine than the D1b_gf strain under similar media conditions. Glutamate cellular pools quantified by LC-MS unveiled distinct trends for growth and indigoidine precursor accumulation (**Figure 4E**). In the strain D1b_gf the increase in glutamate pools were observed between 12 h and 24 h. This increase coincided well with the growth dynamics (**Figure 4E**) and all three substrate consumption profiles (**Figure 4F**). In contrast for strain D1b_gf ΔPP_0897, the glutamate and glutamine pools started to accumulate after 48 h of cultivation which coincided with significant increase in growth post 48 h and consumption profiles of malate and alanine, but not *p*-CA. This result suggests that the cellular production of glutamine/glutamate is strongly linked to the strain growth.

We also confirmed a role for fumarate hydratases in nucleotide biosynthesis. Both the single ΔPP_0897 mutant as well as the D1b_gf ΔPP_0897 strains are resistant to hydroxyurea in rich nutrient growth conditions (**Supplementary Figure 7**). Hydroxyurea is a bacterial anti-metabolite that impairs nucleotide synthesis, suggesting ΔPP_0897 strains are more competent to utilize nucleotide scavenging pathways compared to WT (27,28). The single mutant ΔPP_0897 strain should have a partially active TCA cycle with two other active fumarate hydratase isomers, implying that PP_0897 has a dominant role in maintaining the cellular nucleotide pool over its isomers. This result solidifies that PP_0897 activity is beyond its canonical role as a fumarate hydratase and hints there may be additional activities important when *p-*CA is used as the sole carbon source.

To understand the global changes as a result of the additional deletion of PP_0897 in addition to a medium containing multiple carbon sources, we sampled the two strains at comparable OD_600_ of 0.8 and analyzed the proteomics profiles (**Supplementary Figure 8**). Overall, comparing D1b_gf ΔPP_0897 to D1b_gf in M9 *p*-CA D-alanine L-malate (*p*AM) medium we observed 62 proteins had a significant fold increase (log_2_FC >2.5) and 84 proteins were significantly reduced (log_2_FC <-2.5). Of these 146 proteins, there were 53 proteins associated with metabolic functions including aromatic compound catabolism via beta ketoadipate pathway, TCA, oxidative phosphorylation, branched chain amino acid metabolism. Further 58 proteins that were associated with regulatory functions included several stress proteins, peroxidases and transcriptional regulators. In case of D1b_gf ΔPP_0897 in M9 *p*AM medium compared to M9 D-alanine L-malate medium, we observed 47 proteins had a significant fold increase (log_2_FC >2.5) and 142 proteins were significantly reduced (log_2_FC <-2.5). Of these 189 proteins, there were 62 proteins associated with metabolic functions and 62 associated with regulatory functions. Metabolic proteins were mostly associated with aromatic compound catabolism via beta ketoadipate pathway, oxidative phosphorylation and ABC transport system whereas regulatory proteins were associated with various transcriptional regulators and stress response proteins. Narrowing down to differentially expressed proteins associated with central metabolic pathways with respect to *p*-CA utilization and indigoidine production, we observed that *p*-CA catabolic pathways, ATP synthase and oxidative stress related peroxidases were the most perturbed proteins.

## DISCUSSION

Initially considered to be essential based on results from high-throughput transposon mutagenesis, reports in the literature suggested that PP_0897 could be deleted. For example, deletion analyses of homologous *fumC* fumarate hydratases have been used to probe its function in many processes; a *P. aeruginosa* PAO1 homolog of PP_0897 was deleted and shown to have promiscuous activity on converting mesaconate to (S)-citramalate (29). PP_0897 homologs in other species including *E. coli* and yeast have been reported to indirectly enhance the DNA damage response to double-strand breaks; the metabolite fumarate modulates the bacterial chemotaxis response (30–32). Deletion strains of the eukaryotic yeast homolog of PP_0897, *FUM1*, are viable but growth on non-fermentative carbon sources requires medium supplementation with either aspartic acid, asparagine, or serine (33), parallel with our observations on nutrient auxotrophy in the D1b_gf ΔPP_0897 strain. However, in this study, implementation of this final deletion resulted in a strain that did not grow or produce per the predicted design. While a modulation of the PP_0897 gene enabled systems that recovered growth and production phenotypes, some aspects of the results require additional studies for a better mechanistic understanding. Measurement of glutamine and glutamate levels revealed that glutamate cellular pools could be modulated by changing PP_0897 levels in the D1b_gf strain. However, we did not observe a stoichiometric conversion of these intermediates to the final product indigodine, suggesting cofactor imbalances or other growth and pathway protein expression impacts that remain obscure. In this regard, proteomics data proved instrumental in understanding the strain behavior when PP_0897 activity was altered and could be used to develop context specific models that confirmed the narrow flux required for the FUM reaction. Ultimately, the best experimental solution obtained from the Design 1 strains are those that permitted some activity through this reaction, outperforming strains where the node was fully inactive. Flux through this node is required for biomass formation as well as specific final product titers demanded by the growth coupling design.

The key insight of this study comes from the double sensitivity analysis that enabled evaluation of a single gene intervention representing a limiting enzymatic reaction on both growth and production. Specifically, double sensitivity analysis maps production of the target metabolite and implementation of a gene deletion while optimizing for growth and fitness. In the ideal situation, growth and production are inversely related, where decreasing growth increases the final product. In contrast we find a direct correlation - while intervention of up to three steps yields a configuration with a broad permissible growth and production space, titrating down expression of the last gene, PP_0897, implies that the deletion is inviable because the permissible space is even further constrained. Neither production nor growth can occur. While such a double sensitivity analysis is not a conventional approach used to analyze growth coupled designs, it accurately recapitulates post hoc the experimental evidence in this study. The data-curated model from the promoter titration strain exhibits the overconstrained permissible design space for the growth coupled phenotype in these scenarios.

Advancement in bioproduction and biomanufacturing necessitates the implementation of large sets of gene interventions for strain design and the challenges encountered here may be generally applicable for other final products and hosts. Non-model hosts (e.g. *P. putida* (34), *P. taiwanensis* (35), *Vibrio natriegens* (36)), non-canonical carbon sources (e.g. lignin derived aromatics such as *p*-CA or C_1_ carbon formate) or poorly studied growth formats (e.g. membrane or other alternative bioreactors formats (37,38)) are increasingly found advantageous in these biomanufacturing scenarios. We propose that a stringent threshold for evaluating cutsets may reconcile imperfections in models with their implementation. Enzymatic reactions selected for removal are mapped back to their corresponding coding sequences. Previously, we assumed reactions in the metabolic model mapped to genes encoding only one biological activity and excluded only those that were potentially essential. While we now know that fumarate hydratase PP_0897 is not an essential gene in *P. putida*, its activity was still required for alternative functions when using nonconventional carbon streams like *p-*CA as a carbon source. As such cutsets should be filtered to remove essential genes, known multi-functional enzymes, and sequences where multiple proteins are encoded in the same DNA sequence in a sense-antisense configuration (39). An analysis of available RB-TnSeq (40) datasets indicates several other TCA cycle enzymes may result in a dependency similar to that observed for PP_0897; while transposon mutants are absent from *P. putida*, transposon mutants were recovered in homologs in related *Pseudomonads* and have strong fitness defects across diverse carbon substrates. It is likely that attempts to fully inactivate the citrate synthase or succinate dehydrogenase, for example, would also be difficult to include. These additional precautions will preserve cutset fidelity as enzymatic reactions are translated to encoded genes selected for strain engineering. Algorithms that output potentially over-constrained solutions via strong growth coupling are a powerful approach to obtain strong production phenotypes that are able to meet the stringent titer, rate, and yield demands necessary for an economically viable bioconversion process using renewable carbon streams (41). These approaches will become more valuable as additional catabolic profiles are discovered and introduced into host platforms (35,42,43). Modifying the current growth coupling workflow that accounts for these additional findings in conjunction with functional genomics and ALE could enable streamlined strains for biomanufacturing.

## MATERIALS AND METHODS

### Cultivation of *Pseudomonas putida*

Strains used in this study were propagated using conventional laboratory cultivation protocols for *P. putida* KT2440 (44–46). Engineered strains were grown in a modified M9 minimal medium as previously described (10), and *p*-CA (Sigma-Aldrich, Product No. C9008), L-malic acid sodium salt (Sigma-Aldrich, Product No. M1125) and D-alanine (Sigma-Aldrich, Product No. A7377) were used at the concentrations indicated in the figure legends. All analytes were prepared at a concentration of 0.5 M using reverse osmosis water and filter-sterilized (0.2 mm SFC, Nalgene, Cat. No. 291–4520). The pH of L-malic acid and *p*-CA was adjusted to 6 and 8, respectively, using NaOH pellets (Sigma-Aldrich, Product No. S8045) before filter-sterilization. Strains and plasmids used in this study are described in **Supplementary Table 1 and Supplementary Table 2**.

For production runs, strains were prepared for growth in M9 minimal salt medium with two adaptations as described in (10). 30 mM 3-(N-Morpholino)propanesulfonic acid (MOPS) (Sigma-Aldrich, Product No. 69947) was used as a buffering agent at pH at 7 during the experimental time course. L-arabinose (Sigma-Aldrich, Product No. A3256) was added to induce genes under the *BAD* promoter at a concentration of 1.5 w/v % unless otherwise indicated. Transformation of plasmids and oligos in *P. putida* strains was conducted using standard electroporation techniques described in (7), and strains harboring the overexpression plasmids were kept for no more than two weeks before regeneration. Kanamycin (50 µg/mL) or gentamicin (30 µg/mL) was supplemented to the culture medium as indicated in the respective figures when required.

### Construction of plasmids and targeted genomic mutants via recombineering

In-frame genomic deletions and promoter substitutions were generated exactly as described in (45). All recombineering 90-mer ssDNA oligos and gRNA targeting sequences were synthesized by IDT (Integrated DNA Technologies, Redwood City, CA) and are described in **Supplementary Table 3**. Clones were genotyped with outside-outside colony PCR with NEB OneTaq (New England Biolabs (NEB), Product No. M0482L) following incubation in 20 mM NaOH at 94 °C for 30 min. Synthetic promoter substitutions were verified by sequencing the targeted locus (Azenta Life Sciences, Burlington, MA).

All DNA constructs and oligos were designed using Snapgene (BioMatters Ltd). PCR was conducted using Q5 polymerase (NEB) following manufacturer’s guidelines. Plasmids were assembled using isothermal HiFi assembly (New England Biolabs (NEB), Ipswitch, MA) following manufacturer’s guidelines. Whole plasmid sequencing (Primordium Labs, Monrovia, CA) was used to verify constructs as necessary.

### Indigoidine colorimetric quantification and specific indigoidine productivity assessment

Indigoidine extraction and measurement was described previously in (10) and modified slightly. Briefly, the pH of the culture was adjusted to 7 before sampling if necessary. Samples were then collected and centrifuged at 6,010 x *g* for 3 minutes to precipitate the insoluble indigoidine fraction and cell debris. The cell pellet was resuspended in 100% DMSO pipetting up and down thoroughly, and the mixture was vortexed for 45 min in a Plate Incubator (Benchmark Scientific, Product. No. H6004). Solubilized indigoidine was then separated from the cell debris by centrifugation at 21,130 x g for 1.5 min and its concentration was measured by detecting its absorbance at 612 nm using a microplate reader and converted to g/L by applying the standard curve previously generated with purified indigoidine (10). This study was conducted concurrently alongside Eng and Banerjee 2023 (10) and shares the same base strains and purified indigoidine used for the standard curve generated using *P. putida* fed *p-*CA.

Strains D1b_gf, D1b_gf P_J23109_-PP_0897 and D1b_gf P_PP_0415_-PP_0897 were tested for indigoidine production in M9 60 mM *p*-CA as described above. 24 h post-inoculation, the concentration of the synthesized indigoidine was measured. To calculate specific indigoidine titers (mg/CFU), cultures were diluted 50,000-fold and plated on LB agar to allow colony formation. Colonies were counted 24 hours post incubation at 30 °C. The indigoidine titer (mg/L) was normalized to the total number of viable cells (CFUs/L).

### Glutamine, glutamate, *p*-CA, L-malate and D-alanine quantification by UV-vis HPLC and LC-MS

Culture samples were taken and centrifuged at 6,010 x *g* for 3 minutes at the designated timepoints and supernatants were harvested and frozen at -20 °C until their analysis to quantify the substrate consumption of the engineered strains to calculate the indigoidine production yields (g/g).

When *p*-CA was used as the sole carbon source, its concentration in extracellular supernatant samples was analyzed by HPLC as previously described (10,47). Supernatant samples were defrosted at room temperature and diluted in mobile phase, a solution of 10 mM Ammonium Acetate (Sigma-Aldrich, Cat. No A1542) and 0.07% Formic Acid (Sigma-Aldrich, Cat. No F0507) in HPLC-grade water (Honeywell, Cat. No LC365). A Diode Array Detector (G4212, Agilent Technologies) was used to measure UV absorption at 254 nm, 310 nm, and 280 nm with a Eclipse Plus Phenyl-Hexyl column (250 mm length, 4.6 mm diameter, 5 μm particle size, Agilent Technologies). The flow rate was set to 0.5 mL/min for a 70/30 combination of the mobile phase in water and acetonitrile (Sigma-Aldrich, Cat. No 34851), respectively.

LC-MS was used to monitor substrate co-utilization in M9 minimal media containing *p*-CA, D-alanine and L-malic acid. Supernatants were obtained and preserved as indicated above and the analytes were quenched with 100% methanol (Sigma-Aldrich, Cat. No 34860) and diluted to a final concentration of 50% methanol. For the quantification of glutamine/glutamate pools, both the supernatant and the cell pellet were quenched together with methanol and also analyzed by LC-MS. Due to the spontaneous conversion of glutamine to glutamate at physiological pH values (48,49), we report a summed value of both glutamine and glutamate due to the potential for intracellular spontaneous conversion of glutamine back into glutamate.

LC-MS analysis of these analytes was conducted on a Waters Acquity UPLC BEH Amide column, (100-mm length, 2.1-mm internal diameter, and 1.7-μm particle size; Waters Corporation, Milford, MA, USA) using a 1290 Infinity II UHPLC system (Agilent Technologies, Santa Clara, CA, USA). A sample injection volume of 1 μL was used throughout. The sample tray and column compartment were set to 4 °C and 30 °C, respectively. The mobile phase was composed of 10 mM ammonium acetate, 0.2 % ammonium hydroxide (Sigma-Aldrich), and 5 µM medronic acid (Sigma-Aldrich) in water (solvent A) and 10 mM ammonium acetate, 0.2 % ammonium hydroxide (Sigma-Aldrich), and 5 µM medronic acid (Sigma-Aldrich) in 80 % acetonitrile and 20 % water (solvent B). The solvents were of LC-MS grade and were purchased from HoneyWell Burdick & Jackson (Charlotte, NC, USA). Analytes were eluted via the following gradient conditions: held at 100 %B for 0.5 min, linearly decreased from 100 %B to 75 %B in 2.43 min, linearly decreased from 75 %B to 70 %B in 0.35 min, held at 70 % B for 0.55 min, linearly decreased from 70 %B to 50 %B in 0.2 min, held at 50 %B for 0.3 min, linearly increased to 100 %B in 0.2 min, and held at 100 %B for 2.47 min. The flow rate was held at 0.36 mL/min for 4.33 min, linearly increased from 0.36 mL/min to 0.5 mL/min in 0.2 min, and held at 0.5 mL/min for 2.47 min. The total UHPLC run time was 7 min. The UHPLC system was coupled to an Agilent Technologies 6545 quadrupole time-of-flight mass spectrometer (for LC-QTOF-MS). The QTOF-MS was tuned with Agilent Technologies ESI-L Low concentration tuning mix (at a tenth of its concentration) in the range of 50-1700 m/z. Electrospray ionization via the Agilent JetStream Source (AJS) was conducted in the negative ion mode (for [M - H]-ions) and a capillary voltage of 3500 V was utilized. Drying and nebulizer gases were set to 10 L/min and 20 lb/in2, respectively, and a drying-gas temperature of 300 °C was used throughout. AJS sheath gas temperature and sheath gas flow rate were set to 350 °C and 12 L/min, respectively, while the nozzle voltage was set to 2000 V. The fragmentor, skimmer, and OCT 1 RF Vpp voltages were set to 100 V, 50 V, and 300 V, respectively. The data acquisition rate was set to 0.86 spectra/s. The data acquisition range was from 60-1100 m/z. Data acquisition (Workstation B.08.00) and processing (Qualitative Analysis B.06.00 and Profinder B.08.00) were conducted via Agilent Technologies MassHunter software. Glutamate was quantified via nine-point calibration curve from 0.39 to 100 μM whereas malate, *p*-CA and alanine were quantified via seven-point calibration curves from 0.78 to 50 μM with R2 coefficients of ≥ 0.99.

### BIOLOG Phenotype Microarray

A BIOLOG™ phenotype microarray assay (Cat. No. 12111, & Cat. No. 12112, Hayward CA) was conducted using strain D1b_gf ΔPP_0897. These plates use a tetrazolium dye to detect respiratory products rather than bulk changes in biomass formation as a proxy for cellular growth (50,51) and have been used to characterize various *P. putida* strains (24,25,52,53). A freshly grown single colony was isolated from an LB agar streakout plate after overnight growth from cryostorage and used to inoculate an overnight liquid LB culture at 30 °C. A 1.5 mL aliquot was harvested by centrifugation and washed once with 500 *µ*L M9 medium to remove trace media. Phenotyping plates were prepared exactly as described using the manufacturer’s protocol and incubated at 30 °C without shaking. Redox dye formation was monitored at OD_595_ 24 h and 48 h post inoculation using a Filtermax F5 plate reader (Molecular Devices LLC, San Jose, CA).

### Shotgun Proteomics Analysis

The D1b_gf strains designed in this study were grown in triplicates in M9 60 mM *p*-CA, or M9 50 mM *p*-CA supplemented with 70 mM D-alanine and 70 mM L-malate when indicated, using 10 mL culture tubes. The strains were back-diluted to a starting OD_600_ of 0.05 from a saturated culture, and samples were harvested when cells reached mid-log phase (OD_600_ = 0.8-1) and stored at -80 °C until sample preparation. After all samples were collected, protein was extracted from harvested *P. putida* strain cultures and tryptic peptides were prepared by following established proteomic sample preparation procedures (54). Briefly, cell pellets were resuspended in Qiagen P2 Lysis Buffer (Qiagen Sciences, Germantown, MD, Cat. 19052) for cell lysis. Proteins were precipitated with addition of 1 mM NaCl and 4 x volume acetone, followed by two additional washes with 80% acetone in water. The recovered protein pellet was homogenized by pipetting mixing with 100 mM ammonium bicarbonate in 20% methanol. Protein concentration was determined by the DC protein assay (BioRad Inc, Hercules, CA). Protein reduction was accomplished using 5 mM tris 2-(carboxyethyl)phosphine (TCEP) for 30 min at room temperature, and alkylation was performed with 10 mM iodoacetamide (IAM; final concentration) for 30 min at room temperature in the dark. Overnight digestion with trypsin was accomplished with a 1:50 trypsin:total protein ratio. The resulting peptide samples were analyzed on an Agilent 1290 UHPLC system coupled to a Thermo Scientific Orbitrap Exploris 480 mass spectrometer for discovery proteomics (55). Briefly, peptide samples were loaded onto an Ascentis® ES-C18 Column (Sigma–Aldrich, St. Louis, MO) and separated with a 10 min LC gradient (10% Buffer A (0.1% formic acid (FA) in water) – 35% Buffer B (0.1% FA in acetonitrile)). Eluting peptides were introduced to the mass spectrometer operating in positive-ion mode and were measured in data-independent acquisition (DIA) mode with a duty cycle of 3 survey scans from m/z 380 to m/z 985 and 45 MS2 scans with precursor isolation width of 13.5 m/z to cover the mass range. DIA raw data files were analyzed by an integrated software suite DIA-NN (55). The database used in the DIA-NN search (library-free mode) is the latest Uniprot *P. putida* KT2440 proteome FASTA sequence plus the protein sequences of heterogeneous pathway genes and common proteomic contaminants. DIA-NN determines mass tolerances automatically based on first pass analysis of the samples with automated determination of optimal mass accuracies. The retention time extraction window was determined individually for all MS runs analyzed via the automated optimization procedure implemented in DIA-NN. Protein inference was enabled, and the quantification strategy was set to Robust LC = High Accuracy. Output main DIA-NN reports were filtered with a global FDR = 0.01 on both the precursor level and protein group level. A jupyter notebook written in Python executed label-free quantification (LFQ) data analysis on the DIA-NN peptide quantification report, and the details of the analysis were described in the established protocol (56). Differentially expressed proteins were presented as proteins having absolute log_2_ fold change > 1 and p value ≤ 0.03.

### Constraint-Based Modeling and Simulations

The *P. putida* KT2440 genome scale metabolic model (GSMM) iJN1463 (57) and constraint-based modeling methods (58) were used to simulate the experimental scenarios. Aerobic conditions with *para*-coumarate (*p*-CA), which is represented as Trans-4-Hydroxycinnamate (T4hcinnm) in the GSMM, was used as the sole carbon source to model growth in *p*-CA minimal medium conditions. The lower flux bounds for ATP maintenance demand and *p*-CA exchange were 0.97 mmol ATP/gDCW/h and -10 mmol *p*-CA/gDCW/h, respectively. Flux Balance Analysis (FBA) (59) was used to validate the BIOLOG™ phenotypes observed for the engineered strain by customizing the GSMM to best represent the engineered strain. Excretion of byproducts was initially set to zero, except for the reported secreted products specific to *P. putida* (gluconate, 2-ketogluconate, 3-oxoadipate, catechol, lactate, ethanol, methanol, CO_2_, and acetate) that had an upper flux bound of 1000 mmol/gDCW/h. The heterologous two gene indigoidine production pathway was also added to the GSMM and all four Design One cutset gene targets (PP_0944, PP_1755, PP_1378 and PP_0897) were deleted. For each of the BIOLOG™ carbon sources, an uptake flux of -10 mmol/gDCW/hr was set for the respective exchange reaction and we used FBA to maximize the biomass reaction, which was selected as the objective function. The production envelope was generated using robustness analysis using the Design One cutset configured GSMM and *p*-CA as the sole carbon source, biomass reaction as the objective function and indigoidine as the control reaction (20,21). For the glutamine production simulations, pseudo reactions for external export (DF_gln L) and exchange (EX_gln L_e) were added to the model and the EX_gln L_e reaction was used as the objective when simulating for glutamine production.

Double Robustness analysis (20,21) was performed to check for sensitivity of biomass objective function towards the change in flux magnitude through a pair of reactions, i.e. glutamine or indigoidine biosynthesis reaction (EX_gln L_e or EX_ind_e) and the fumarase reaction (FUM). COBRA Toolbox v.3.0 (58) in MATLAB R2017b (Mathworks Inc., Natick, MA, USA) was used for FBA and Double Robustness analysis simulations with Gurobi Optimizer 8.1 (http://www.gurobi.com/) as the solver.

### Proteomics data integration to create context specific models

The proteomics data and the *P. putida* D1b_gf reduced model, iMATD1b539, were used to extract context specific models for the two promoter based strains. We used the integrated metabolic analysis tool (iMAT) algorithm (60), which uses discrete levels for expression values: low, medium and high. iMAT allowed the integration of proteomics data into the GSMM and maximized highly and minimized lowly expressed reactions from the model. We used the log_2_ fold changes for protein counts with respect to D1b_gf in P_J23109_ and P_PP_0415_ strains to perform this classification with a cutoff of half a standard deviation above the mean of the log_2_FC values, for active reactions, and half a standard deviation below for inactive reactions. We used the iMAT algorithm to iteratively search for a set of active reactions, within the cutoff range, linked to the levels of the associated proteins that simultaneously are able to result in biomass and indigoidine production. Simulations were performed using the iMAT implementation in the Cobra Toolbox with Matlab 2017b (Mathworks Inc., Natick, MA, USA) and CPLEX 12.8 as the solver.

## Author Contributions

DB, TE and AM conceptualized the work. DB developed and implemented the computational method. JM and TE generated strains and executed the molecular biology and growth analysis experiments. YC, JWG and CJP procured and analyzed the proteomics data. EEKB procured and analyzed the metabolomics data. DB, JM, TE and AM interpreted the results. DB and TE wrote the original draft. DB, JM, TE and AM were involved in writing, reviewing and editing the figures and manuscript. AM and TE supervised the study. AM acquired funding. All authors read and approved the final manuscript.

## Acknowledgements

We thank Jeff Czajka (PNNL) for insightful comments on this manuscript. This material is based upon work supported by the Joint BioEnergy Institute, U.S. Department of Energy, Office of Science, Biological and Environmental Research Program under Award Number DE-AC02-05CH11231 with Lawrence Berkeley National Laboratory. The funder played no role in study design, data collection, analysis and interpretation of data, or the writing of this manuscript.

## Data Availability

The generated mass spectrometry proteomics data have been deposited to the ProteomeXchange Consortium via the PRIDE partner repository (61). Data and additional information required to reanalyze the data reported in this work paper is available from the lead contact upon request.

All strains and plasmids used in this study are described in **Supplementary Table 1** and **Supplementary Table 2** and are available upon request.

