## Supplementary Information for "Bottlenecks in the Implementation of Genome Scale Metabolic Model Based Designs for Bioproduction from Aromatic Carbon Sources"

**Supplementary Table 1. Strains Used in This Study.**

| Strain ID | Genotype | Relevant Figure(s) | Reference |
|---|---|---|---|
| JBEI_228,473 | *P. putida* KT2440 PP_5402-intergenic::$P_{BAD}$-*Sc.bpsA,Bc.sfp* | 2,SF1, SF3,SF7 | Banerjee and Eng, 2020 |
| JBEI_256,060 | *P. putida* KT2440 PP_5402-intergenic::$P_{BAD}$-*Sc.bpsA,Bc.sfp* ΔPP_0897 | 2,SF7 | This study |
| JBEI_233,088 | *P. putida* KT2440 PP_5402-intergenic::$P_{BAD}$-*Sc.bpsA,Bc.sfp,Ps.glnA* ΔPP_1378 ΔPP_1755 ΔPP_0944 ΔfleQ/ΔPP_4373 serial passaging via ALE (“D1b_gf”) | 1,2,4,SF1, SF2,SF3, SF6,SF7, SF8 | Eng and Banerjee, 2023 |

| | | | |
|---|---|---|---|
| JBEI_256,069 | *P. putida* KT2440 *PP_5402-intergenic::$P_{BAD}$-Sc.bpsA,Bc.sfp,Ps.glnA* ΔPP_1378 ΔPP_1755 ΔPP_0944 Δ*fleQ*/ΔPP_4373 serial passaging via ALE ΔPP_0897 (D1b_gf ΔPP_0897) | 1,2,4,SF2, SF5,SF6, SF7,SF8 | This study |
| JBEI_236,231 | *P. putida* KT2440 *PP_5402-intergenic::$P_{BAD}$-Sc.bpsA,Bc.sfp,Ps.glnA* ΔPP_1378 ΔPP_1755 ΔPP_0944 ΔfleQ/ΔPP_4373 serial passaging via ALE $P_{J23109}$-PP_0897 | 2,SF1,SF 3 | This study |
| JBEI_236,232 | *P. putida* KT2440 *PP_5402-intergenic::$P_{BAD}$-Sc.bpsA,Bc.sfp,Ps.glnA* ΔPP_1378 ΔPP_1755 ΔPP_0944 ΔfleQ/ΔPP_4373 serial passaging via ALE $P_{PP\_0415}$-PP_0897 | 2,SF1,SF 3 | This study |
| JBEI_256,070 | *E. coli* DH10-β pTE408 {$P_{BAD}$ araC BBR1 gntR} | | This study |
| JBEI_256,073 | *E. coli* DH10-β pTE640 {$P_{BAD}$-PP_0897 araC BBR1 gntR} | | This study |
| JBEI_256,077 | *E. coli* DH10-β pTE612 {PP_0897-*gRNA lacMp-RBSopt-aCpf1 BBR1 kanR/neoR lacI-*} | | This study |
| JBEI_256,078 | *E. coli* DH10-β pTE614 {$P_{PP\_0897}$-gRNA *lacMp-RBSopt-aCpf1 BBR1 kanR/neoR lacI-*} | | This study |

**Supplementary Table 2. Plasmids Used in This Study.**

| **Plasmid name** | ***Miscellaneous Notes*** | **Source** |
|---|---|---|
| pTE640 | $P_{BAD}$-PP_0897 *araC BBR1 gntR* | This study; JBEI_256,094 |
| pTE408 | $P_{BAD}$ *araC BBR1 gntR* (empty vector) | This study; JBEI_256,093 |
| pTE612 | $P_{J23119}$-PP_0897-*gRNA lacMp-RBSopt-aCpf1 BBR1* kanR/neoR LacI- (deletion) | This study; JBEI_256,095 |
| pTE614 | $P_{J23119}$-PP_0897-*gRNA lacMp-RBSopt-aCpf1* BBR1 kanR/neoR LacI- (promoter editing) | This study; JBEI_256,096 |

**Supplementary Table 3. List of recombineering oligos and gRNAs used in this study.**

| Oligo number | Sequence (5' - 3') | Notes |
|---|---|---|
| TEAM-2558 | CGAGCGTCTTTTTGAATGTTCCCGTCTTTAAGAGGAGCGCGCTGCCCTGATGTAACTGCGGCGGCTGCTTCGCGGGCAAGCCCGCTCCCA | Recombineering oligo for precise PP_0897 deletion |
| TEAM-3299 | TTCCGTCGGCCTGACCTGCCGGCGCCTTGCAAGGCGCTTCGACGT CCTTCATCGCTTTCCCTGAGTTACAATGCGCGCCACCGTGAATTTGCCAACAGGTGCGCC atgACCGTGATCAAGCAAGACGACCTGATTCAGAGCGTCGCCGAC | Recombineering oligo to replace PP_0897 promoter with low protein abundance protein PP_0415/PP_0416 promoter |
| TEAM-3300 | TTCCGTCGGCCTGACCTGCCGGCGCCTTGCAAGGCGCTTCGACGT TTACAGCAGCCAGCCAGGGACGGCTAGC atgACCGTGATCAAGCAAGACGACCTGATTCAGAGCGTCGCCGAC | Recombineering oligo to replace PP_0897 promoter with pJ23109 low activity promoter from Anderson collection |
| TEAM-2582 | GCCGCTGGCAATCGATATGAAAC | Primer for PP_0897 colony PCR |
| TEAM-2583 | TGGTGTGTGAGGTAGATTTGGAGG | Primer for PP_0897 colony PCR |
| TEAM-2593 | GTAAGGCACATTCAGCCCCG | Sequencing primer; PP_0897 promoter |
| TEAM-2594 | CCTTGTCGCCGGGGACTATG | Sequencing primer; PP_0897 promoter |
| TEAM-3033 | AATGTTCCCGTCTTTAAGAGG | gRNA sequence targeting PP_0897 promoter (pTE614 aCpf1 assembly) |
| TEAM-2559 | ATCCAGGCCATGCACGAGGCC | gRNA sequence targeting PP_0897 (pTE612 aCpf1 assembly) |

**Supplementary Table 4. List of BIOLOG™ metabolites tested in this study.**

| **Metabolite** | **BIOLOG™ $OD_{595}$** | ***In-silico* Growth** |
|---|---|---|
| L-Arabinose | -0.0025 | 0 |
| N-Acetyl-DGlucosamine | -0.0007 | 0 |
| D-Saccharic Acid | -0.0028 | 0 |
| Succinic Acid | -0.0022 | 0 |
| D-Galactose | -0.0003 | 0 |
| L-Aspartic Acid | -0.004 | 0.472 |
| L-Proline | -0.0029 | 0 |
| D-Alanine | 0.7416 | 0.423 |
| D-Trehalose | -0.0036 | 0 |
| D-Mannose | 0.253 | 0.950 |
| Dulcitol | -0.0023 | 0 |
| D-Serine | -0.0063 | 0.381 |
| D-Sorbitol | -0.0106 | 0 |
| Glycerol | 0.5234 | 0.586 |
| L-Fucose | -0.0058 | 0 |
| D-Glucuronic acid | -0.0077 | 0 |
| D-Gluconic Acid | 0.7312 | 0.916 |
| D,L-α-GlycerolPhosphate | -0.0057 | 0 |
| D-Xylose | 0.0076 | 0 |
| L-Lactic Acid | 0.6597 | 0.386 |
| Formic Acid | 0.036 | 0.070 |
| D-Mannitol | -0.0104 | 0 |
| L-Glutamic Acid | -0.0108 | 0 |
| D-Glucose-6-phosphate | -0.0014 | 0 |
| D-Galactonic Acid-γ-Lactone | -0.0113 | 0 |
| D,L-Malic Acid | 0.5018 | 0.46 |
| D-Ribose | 0.0169 | 0.791 |
| Tween 20 | 0.0521 | 0 |

| | | |
|---|---|---|
| L-Rhamnose | -0.0089 | 0 |
| D-Fructose | 0.5086 | 1.003 |
| Acetic Acid | -0.0029 | 0.194 |
| α-D-Glucose | 0.7413 | 0.962 |
| Maltose | -0.0064 | 0 |
| D-Melibiose | -0.0065 | 0 |
| Thymidine | -0.0083 | 0 |
| L-Asparagine | -0.001 | 0.472 |
| D-Aspartic Acid | -0.0074 | 0 |
| D-Glucosaminic acid | -0.0079 | 0 |
| 1,2-Propanediol | -0.008 | 0 |
| Tween 40 | 0.0237 | 0 |
| α-Keto-Glutaric acid | 0.0012 | 0 |
| α-Keto-Butyric acid | -0.0133 | 0 |
| α-Methyl-D-galactoside | -0.0079 | 0 |
| α-D-Lactose | -0.0073 | 0 |
| Lactulose | 0.0005 | 0 |
| Sucrose | -0.0082 | 0 |
| Uridine | 0.0195 | 1.266 |
| L-Glutamine | -0.0222 | 0 |
| m-Tartaric Acid | -0.0011 | 0 |
| D-Glucose-1-phosphate | -0.009 | 0 |
| D-Fructose-6-phosphate | -0.0044 | 0 |
| Tween 80 | 0.015 | 0 |
| α-Hydroxy Glutaric Acid-γ-Lactone | 0.001 | 0 |
| α-Hydroxy Butyric Acid | -0.0047 | 0 |
| ß-Methyl-DGlucoside | -0.007 | 0 |
| Adonitol | -0.0006 | 0 |
| Maltotriose | 0.0334 | 0 |
| 2-Deoxy Adenosine | 0.0044 | 0 |
| Adenosine | 0.1711 | 1.284 |

| | | |
|---|---|---|
| Glycyl-L-Aspartic acid | -0.0052 | 0 |
| Citric acid | 0.2168 | 0 |
| myo-Inositol | -0.0095 | 0 |
| D-Threonine | -0.0055 | 0 |
| Fumaric Acid | -0.0064 | 0 |
| Bromo Succinic acid | 0.02 | 0 |
| Propionic Acid | -0.0173 | 0 |
| Mucic Acid | 0.0121 | 0 |
| Glycolic Acid | -0.0038 | 0.167 |
| Glyoxylic Acid | -0.0026 | 0 |
| D-Cellobiose | -0.0024 | 0 |
| Inosine | 0.3661 | 1.262 |
| Glycyl-LGlutamic Acid | 0.0124 | 0.094 |
| Tricarballylic acid | -0.0067 | 0 |
| L-Serine | 0.1603 | 0.388 |
| L-Threonine | 0.0034 | 0.591 |
| L-Alanine | 0.6446 | 0.423 |
| L-Alanyl-Glycine | 0.3024 | 0.713 |
| Acetoacetic Acid | 0.0005 | 0.442 |
| N-Acetyl-ß-DMannosamine | -0.0052 | 0 |
| Mono-Methyl succinate | -0.0067 | 0 |
| Methyl Pyruvate | 0.1388 | 0 |
| D-Malic acid | 0.0007 | 0 |
| L-Malic Acid | 0.6336 | 0.46 |
| Glycyl-L-Proline | 0.0176 | 0 |
| p-Hydroxy Phenyl Acetic Acid | 0.0079 | 0 |
| m-Hydroxy Phenyl Acetic Acid | 0.0026 | 0 |
| Tyramine | 0.0021 | 0 |
| D-Psicose | 0.004 | 0 |
| L-Lyxose | -0.0039 | 0 |
| Glucuronamide | 0.0034 | 0 |

| | | |
|---|---|---|
| Pyruvic Acid | 0.182 | 0.335 |
| L-Galactonic Acid-γ-Lactone | 0.0004 | 0 |
| D-Galacturonic acid | 0.0004 | 0 |
| Phenylethyl-amine | 0.0049 | 0.284 |
| 2-Aminoethanol | 0.0043 | 0.318 |

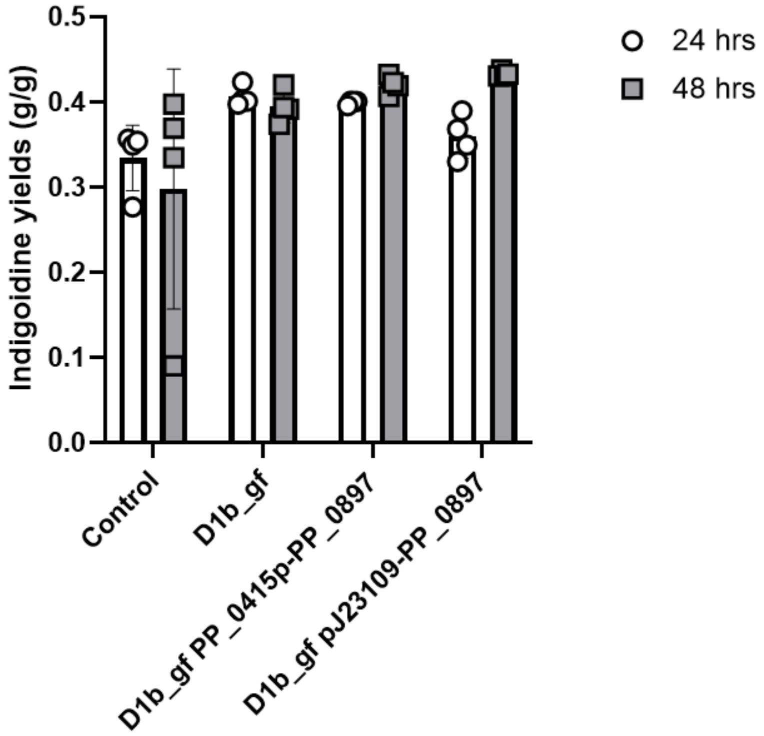


**Supplementary Figure 1. Indigoidine production yields in PP_0897 promoter variants.** Strains were adapted in M9 60 mM *p*-CA and then were prepared for a production run in 24 well deep well plates and induced with 1.5% (w/v) arabinose. The relevant genotype of each strain is indicated below each bar. Indigoidine yields are reported as grams of indigoidine per gram of *p*-CA (g/g). Error bars represent mean ± S.D. (n=4).

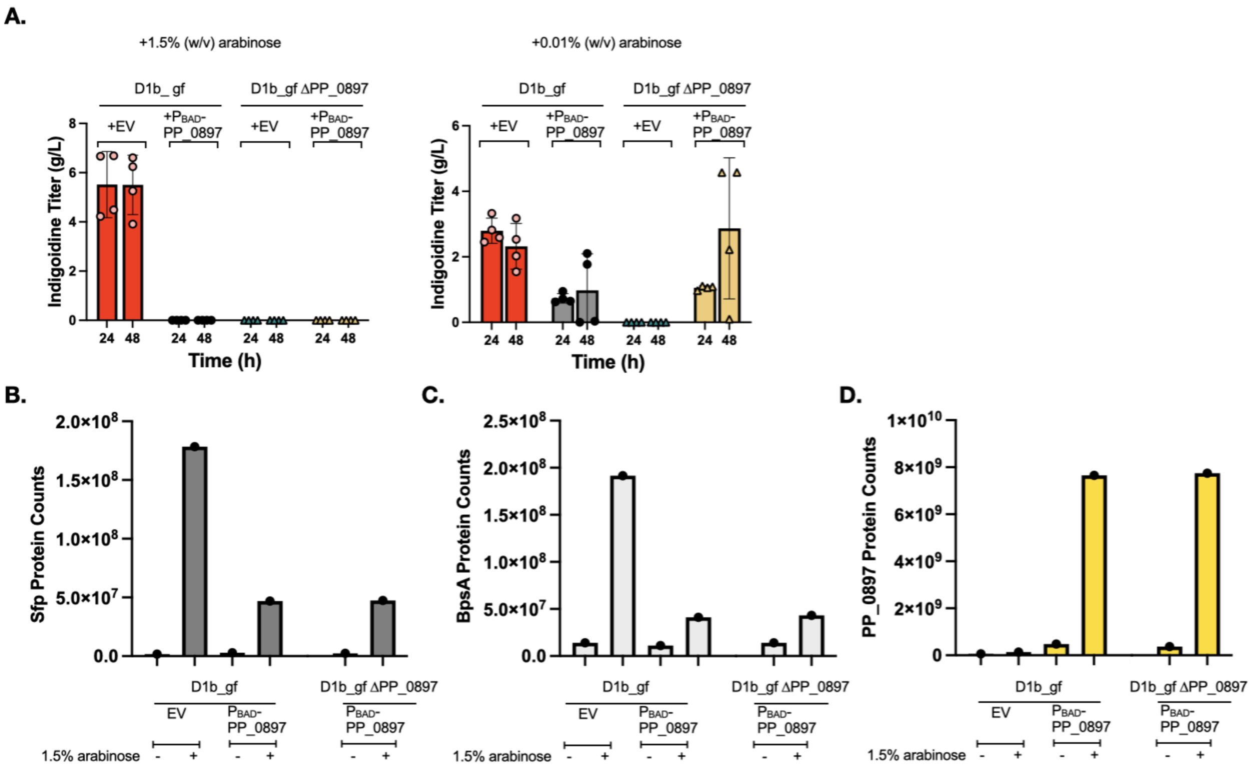


**Supplementary Figure 2. Supporting information related to PP_0897 overexpression.** D1b_gf and D1b_gf ΔPP_0897 were transformed with either an empty vector plasmid (pTE408) or a $P_{BAD}$-PP_0897 overexpression construct. (A) Indigoidine production (refer to Figure 3) with either 0.01 % or 1.5 % (w/v) arabinose added to the media. (B-D) Proteomics analysis of indigoidine pathway proteins Sfp and BpsA as well as PP_0897 protein counts. The indicated strains were grown with or without 1.5 % (w/v) arabinose to induce the *BAD* promoter and allowed to grow at 30 ˚C for an additional 5 hours before samples before proteomics analysis. The relevant strain genotype and presence or absence of arabinose is indicated in the sample legend below the graph. Error bars represent mean ± S.D. (n=4) in panel A.

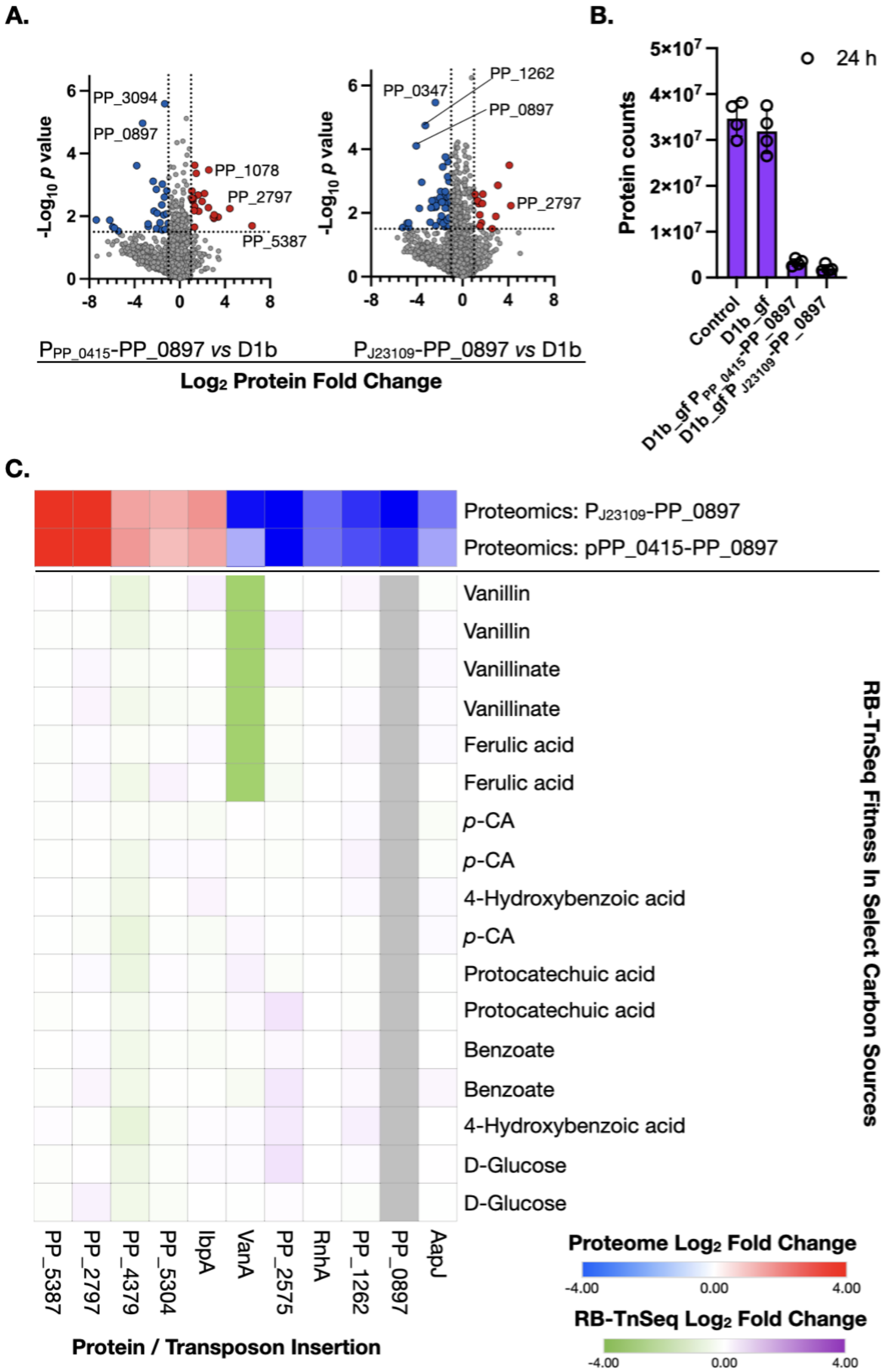


**Supplementary Figure 3: Omics analysis** (A) Global protein changes in the D1b_gf strains harboring PP_0897 promoter variants. The shotgun proteomic dataset for the indicated PP_0897 promoter variants are plotted as volcano plots comparing differential protein counts between the promoter variant strain and the parental D1b_gf strain. (B) PP_0897 protein levels of the indicated strain genotypes were assayed using LC-MS/MS shotgun proteomics. The control strain is a WT *P. putida* KT2440 strain harboring the indigoidine cassette. Proteomics analysis was done using four biological replicates. (C) RB-TnSeq fitness profiling of differentially changed proteins common between both PP_0897 promoter mutants. Error bars represent mean ± S.D. (n=4) in B.

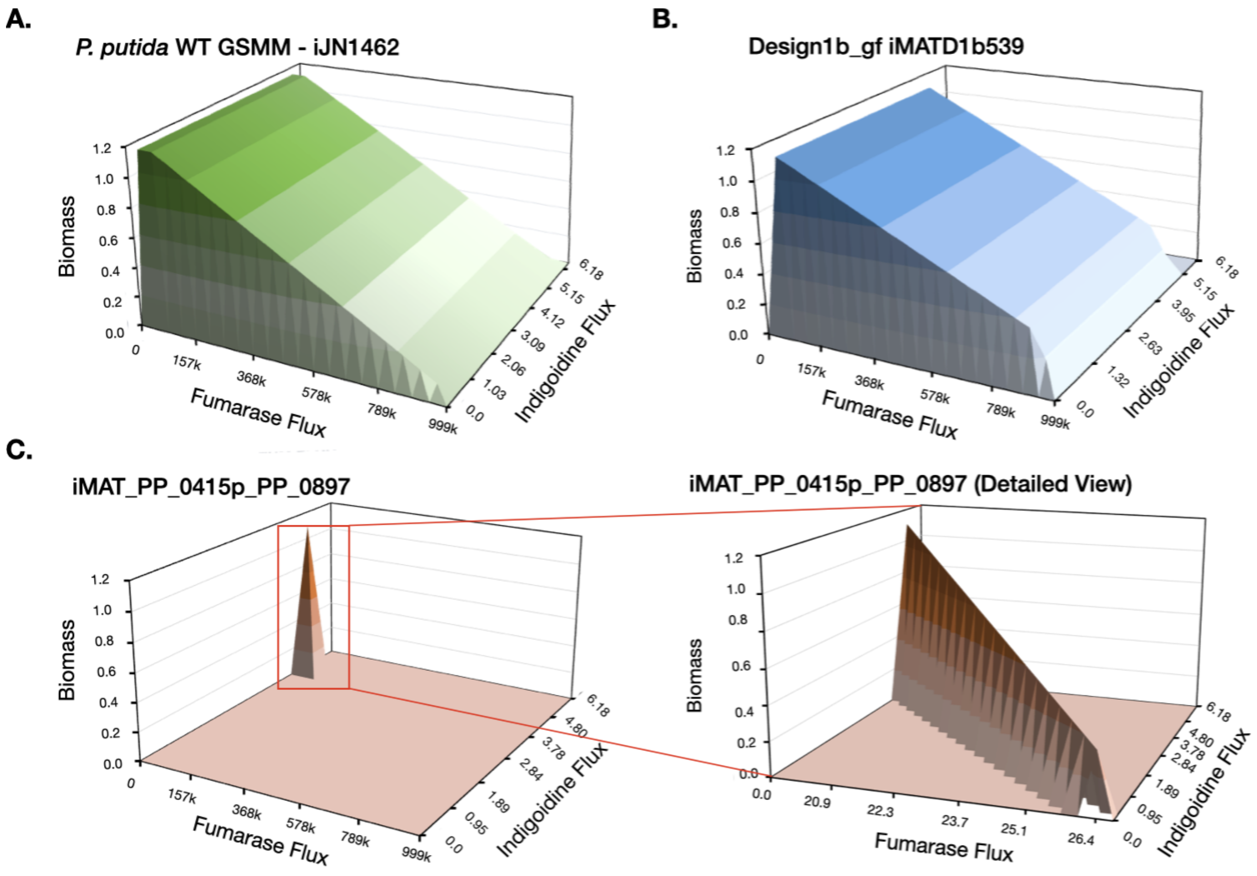


**Supplementary Figure 4. Analysis of proteomics-constrained context-specific metabolic models shows that a narrow permissible fumarase flux space exists for growth coupled indigoidine production scenarios.** A double robustness analysis is applied to determine the tradeoff between growth, fumarase flux and indigoidine formation. Double robustness analysis 3D plots of the genome scale metabolic model (GSMM) of *P. putida* WT (iJN1462), (A), context specific D1b_gf reduced model (B) and (C) Context specific D1b_gf PP_0415p-PP_0897 model (iMAT_PP_0415p_PP_0897). A red box indicates the region of interest replotted in the detailed view.

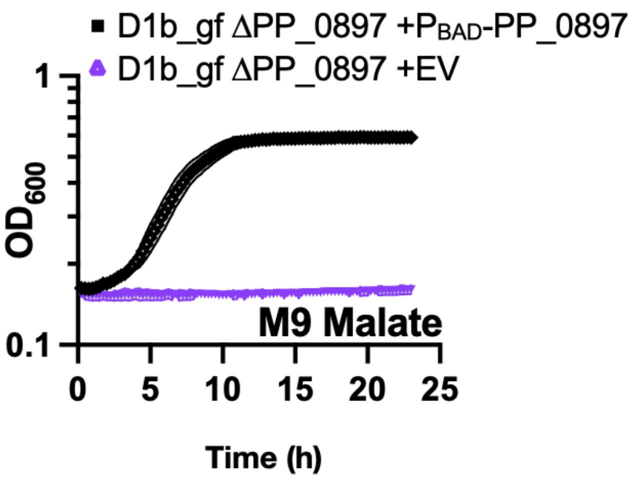


**Supplementary Figure 5.** 70 mM malate supported growth as the sole carbon source in control strains, indicating malate transport was not a limiting factor. Shaded area represents mean ± S.D. (n=3).

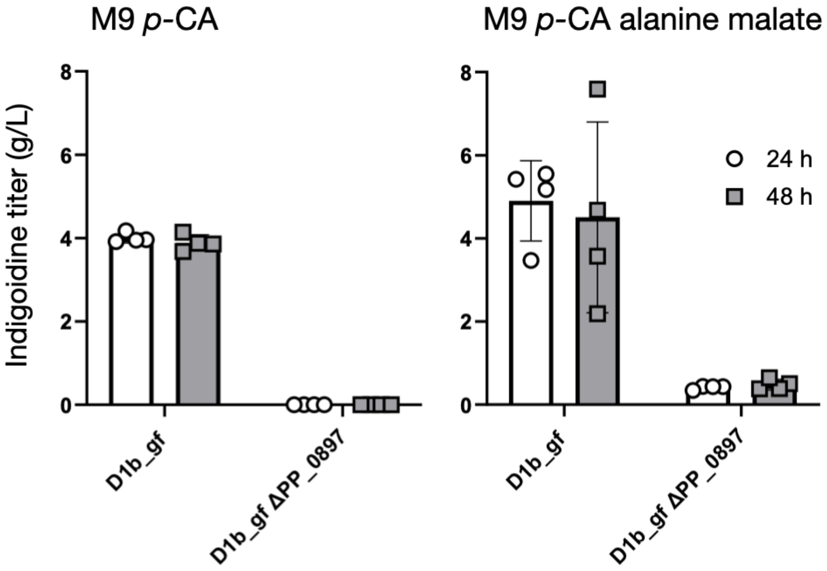


**Supplemental Figure 6. Indigoidine production in strains D1b_gf and D1b_gf ΔPP_0897 in M9 60 mM *p*-CA medium (left) and M9 50 mM *p*-CA, 70 mM D-alanine and 70 mM L-malate medium (right).** Strains were adapted two times in M9 minimal medium for production as described (see materials and methods) using M9 *p*-CA when the production run was done in the same medium, and M9 alanine when the production run was performed in M9 *p*-CA, alanine, malate medium. These experiments were performed in quadruples in 24-deep-well plates using a fill volume of 1.5 mL and 1.5 % of arabinose as the production inducer. Error bars represent mean ± S.D. (n=4).

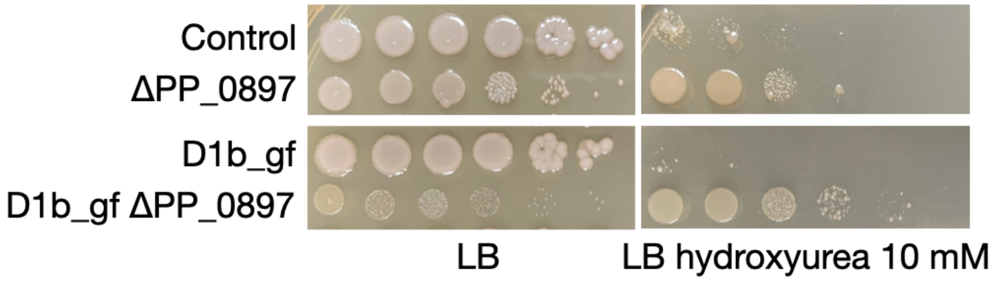


**Supplemental Figure 7**. **PP_0897 deletion strains are resistant to hydroxyurea.** Strains of the indicated genotype were serially diluted onto solid agar plates supplemented with LB and 10 mM hydroxyurea as indicated. Plates were incubated at 30 ˚C and inspected every 24 hours to assess growth. The LB plates were photographed 48 hours post incubation and the LB hydroxyurea plates were photographed 72 hours post incubation.

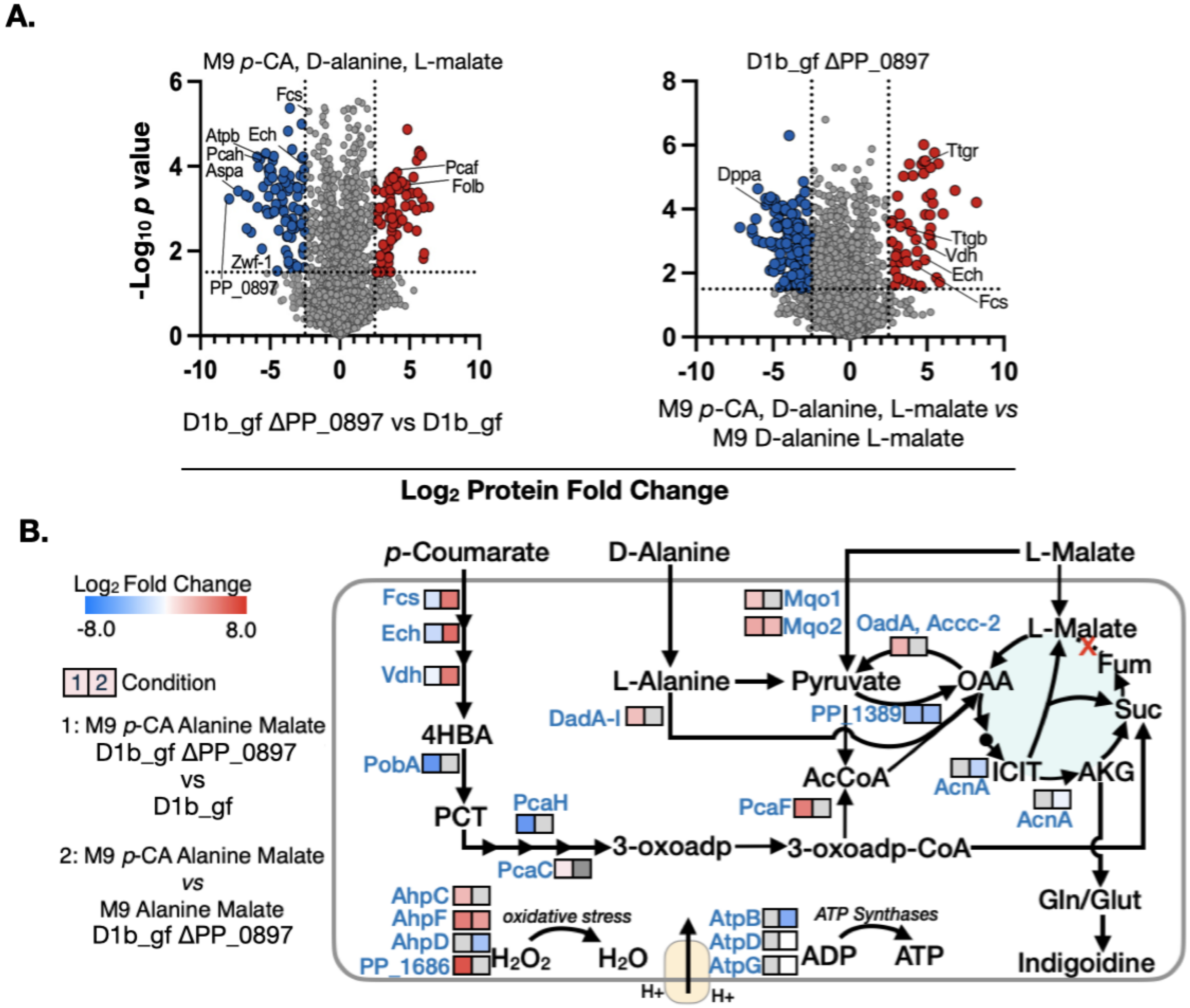


**Supplementary Figure 8. *p*-CA catabolic pathway proteins are downregulated when all fumarate hydratases are deleted.** (A) Volcano plots showing the global protein changes in the D1b_gf ΔPP_0897 strain compared to D1b_gf strain in multiple carbon source media conditions. The shotgun proteomic dataset for the indicated strains and media conditions are plotted as volcano plots comparing differential protein counts between the different strains in PAM medium as well as D1b_gf strain in different media compositions. (B) Differential protein expression counts overlaid onto a metabolic map. Abbreviations used - 4HBA: 4-hydroxybenzoate; Fum: Fumarate; Glu: Glutamate; Gln: Glutamine; ISO: Isocitrate; OAA: Oxaloacetate; PCT: Protocatechuate; *p*-CA: *para*-coumarate; Suc: Succinate.

### Supplementary Results for Model Validation using BIOLOG™ Phenotypic Data

For the PM1 BIOLOG™ phenotype microarray plate, 15 false predictions (2 false positives,13 false negatives) may be a result of missing gaps or redundancy in the GSMM. Of the 13 false positives, in the case of formic acid and glycyl-L-glutamic acid, the predicted biomass was less than 0.1. For acetic acid, glycolic acid, phenylethyl-amine and 2-aminoethanol, the predicted biomass was in the range of 0.1 to 0.32 after 24 h and were not suitable for measurements using BIOLOG$^{TM}$ plates. For additional carbon sources including adenosine, L-alanyl-glycine and inosine we were able to predict growth after adding transport reactions to the metabolic model. Of the 2 false negative carbon sources, methyl pyruvate is absent from the GSMM, whereas citrate has no clear explanation, and would require further investigation into the GSMM accuracy. In the case of the PM2A BIOLOG$^{TM}$ plate, there were only two additional carbon sources where growth was observed: dextrin and D, L-carnitine. Dextrin was among the 74 metabolites in the PM2A plate that do not exist in the GSMM. Of the remaining 21 metabolites, 11 were predicted accurately using the GSMM including biomass formation using L-carnitine for the D1b_gf ΔPP_0897 strain. There were 10 metabolites, mostly amino acids, that were predicted as false positives.
